# Simultaneous single-cell calcium imaging of neuronal population activity and brain-wide BOLD fMRI

**DOI:** 10.1101/2023.11.14.566368

**Authors:** Rik L.E.M. Ubaghs, Roman Boehringer, Markus Marks, Helke K. Hesse, Mehmet Fatih Yanik, Valerio Zerbi, Benjamin F. Grewe

## Abstract

Blood Oxygen Level-Dependent (BOLD) functional Magnetic Resonance Imaging (fMRI) allows for non-invasive, indirect recordings of neural activity across the whole brain in both humans and animals. However, the relationship between the local neural population activity and the vascular activity is not completely understood. To investigate this relationship, we present a novel MRI compatible single-photon microscope capable of measuring cellular resolution Ca^2+^ activity of genetically defined neurons during whole-brain BOLD fMRI in awake behaving mice. Using this combined imaging approach, we found a difference in activity patterns between cells which was dependent on their location with respect to the vasculature. Notably, neurons near the vasculature showed pronounced negative activity during contralateral whisker movements at 3 Hz. In a second proof of concept experiment, we demonstrate the potential of recording both local neural activities, like those in the barrel field (SSp-bfd), and BOLD fMRI readings from interlinked brain regions. In sum, the presented technological advancement paves the way for studies examining the interplay between local brain circuits and overarching brain functions. In addition, the new approach enhances our understanding of the vascular BOLD fMRI signal, providing insights into the determinants of local neurovascular functions and the brain’s organizational framework across various scales.

## Main

Brain function emerges from coordinated interactions of global brain areas and local neural circuits. Investigating brain functioning therefore benefits from the availability of experimental methods capable of probing neural activity throughout the whole brain, preferably with a high spatial and temporal resolution at the level of local neuronal circuits.

Widely adopted imaging methods such as functional Magnetic Resonance Imaging (fMRI) allow non-invasive whole brain recordings but compromise spatiotemporal resolution and source specificity for whole-brain coverage and non-invasiveness. For example, Blood Oxygen Level-Dependent (BOLD) fMRI measures neural activity indirectly through changes in blood oxygenation and volume^1,2^, which result from relatively slow processes involved in neurovascular coupling. The BOLD fMRI signal is therefore considered as a slow, mean-proxy of local brain activity, with a limited ability to resolve local neural circuit dynamics^3^.

In contrast, opto-physiological recording methods, such as *in vivo* single- or multi-photon imaging, are capable of capturing the membrane voltages or Ca^2+^ levels of genetically defined neural circuits at cellular resolution^4,5^. Recent Ca^2+^ imaging studies in rodents demonstrated the simultaneous monitoring of thousands of individual cells at kHz sampling rates^6,7^. However, optical methods typically require invasive surgery to gain access to the brain^8,9^, and are only capable of simultaneously measuring up to a few cortical brain regions of interest in animals^10^.

To bridge the gap between local neural circuit activity and the hemodynamic BOLD fMRI signal, researchers have started to combine optical imaging methods with fMRI. A common *in vivo* combined imaging approach (for *in vitro* see 11) has been to perform sequential fMRI and Ca^2+^ imaging within the same subject^12^. While this approach allows to compare the averaged local neural and hemodynamic responses, it is unable to control for variability between trials, sessions, and experimental environments^13,14^.

To better understand the temporal relationship between neural circuit activity and BOLD fMRI, several research groups developed methods for simultaneous fiber photometry-based Ca^2+^ imaging and BOLD fMRI in anesthetized rodents^15–19^. For example, Schulz et al. (2012) used combined fiber-photometry and BOLD fMRI to show that the BOLD fMRI signal reflects not only neural sources, but also relates to local, slow oscillating Ca^2+^ waves in astrocytes. More recently, Lake et al. (2020) expanded on combined fiber-photometry fMRI by developing a multi-fiber bundle to record simultaneous macro-scale Ca^2+^ and BOLD activity across the whole cortex. Their results show that the BOLD fMRI signal across the cortex can be partially reconstructed from local neural activity with high spatial precision, laying the foundation for a better understanding of the neural underpinnings of the BOLD fMRI signal.

These previous studies have demonstrated that combined optical and BOLD fMRI approaches offer a powerful way of exploring neurovascular dynamics. However, fiber-based methods do not provide the spatial resolution needed to distinguish single neurons, and instead group together (non-)neuronal cellular and neuropil (synaptic and dendritic) signals. To overcome this limitation, we developed a novel MRI compatible single-photon microscope capable of measuring cellular resolution Ca^2+^ activity of genetically defined neurons without inducing artifacts during whole-brain BOLD fMRI in awake behaving mice. By offering a cellular resolution readout of neuronal activity within the context of whole-brain functional imaging, our approach for the first time allows researchers to compare the activity of a population of genetically defined neurons to the vascular activity throughout the rodent brain.

## Results

### Combining fMRI and population calcium imaging technologies

Ca^2+^ imaging is typically performed with microscopes that consist of materials and electronics incompatible with the magnetic field of Magnetic Resonance Imaging (MRI) scanners. To combine Ca^2+^ imaging and MRI, we designed a small MRI compatible microscope that functions within the limited space of a standard small-animal MRI volume coil (70 mm diameter; RF RES 300 1H 089/072 QSN TR AD, BRUKER BioSpin MRI GmbH), and is compatible with a surface coil (RF SUC 300 1H LNA AV3, BRUKER BioSpin MRI GmbH; **Fig. 1a**). The main components of our MRI compatible imaging device include a CCD camera (MRC HighResolution, MRC Systems GmbH), fluorescence light excitation and imaging optics, and a custom 3D printed microscope body (**Fig. 1a,b**; github.com/rlemubaghs/open_mrscope.git).

**Figure 1.**
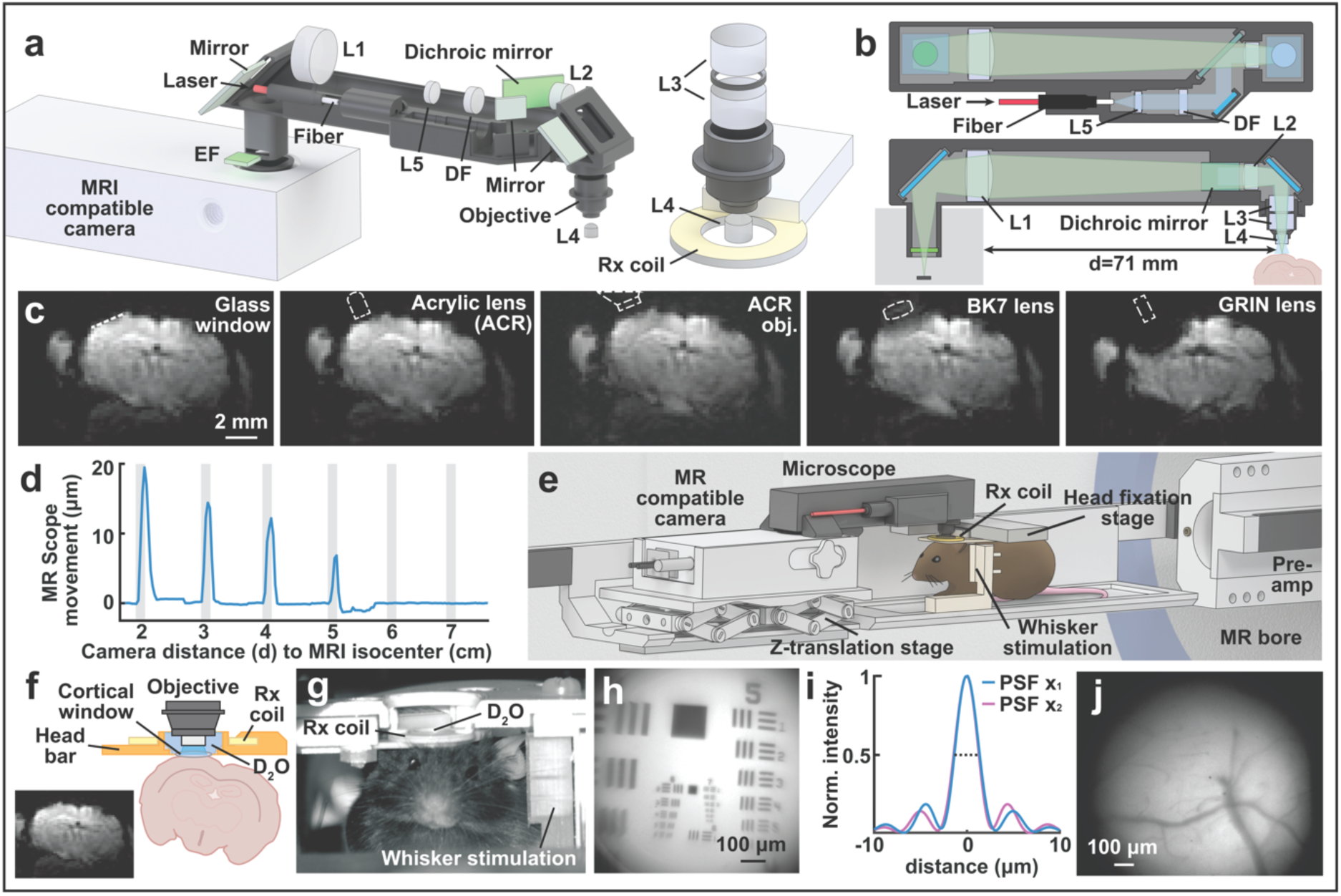
MRI compatible fluorescence microscope design for combined BOLD fMRI and calcium imaging. **a**, CAD rendering of the MRI compatible microscope (left) and objective (right). Excitation light from an off-site laser is collimated with a double-convex lens (L5) and passed through a diffuser (DF). Fluorescent emission is collected by an objective consisting of three small diameter acrylic aspheric lenses (L3, L4). The collected light is relayed through two achromatic lenses (L1, L2), and projected onto the MRI compatible CCD camera through an emission filter (EF). **b,** Schematic of the excitation (blue) and emission (green) optical pathways. The distance between the CCD camera housing and the sample is 71.1 mm. **c,** BOLD EPI of a mouse brain (*in situ*) with and without optical components placed close to the Ca^2+^ imaging site above the barrel cortex. From left to right: glass window only, acrylic lens, acrylic lens and HDT180 objective, BK7 objective lens, GRIN objective lens. **d,** Microscope displacement induced by RF excitation pulses with respect to the distance between the MRI compatible camera and the isocenter of the B0 magnetic field. **e,** Complete MRI imaging setup including the microscope, a translational stage for focusing, the receiver (Rx) coil, the whisker stimulation apparatus, the head fixation stage, the pre-amplifier, and the MRI scanner. **f,** Schematic drawing of the implant, Rx coil, head bar and cranial window. Inset: BOLD EPI image with D_2_O immersion. **g,** Photo of the same implant setup showing a mouse fixed in the behavioral cradle. **h,** Fluorescence images taken with the microscope above the barrel cortex. **i,j,** High resolution image of a microscope calibration slide (**i**) and the simulated optical resolution of the microscope (**j**) with a full-width-half-maximum (FWHM, dashed line) of 2.5 micron.

To minimize artifacts and optimize the signal-to-noise ratio of BOLD fMRI, we took several measures to ensure the homogeneity of the magnetic field and to avoid electromagnetic interference, without compromising fluorescent signal transmission. These include (**1**) shielding or positioning MRI incompatible microscope electronics outside the MRI’s magnetic field, (**2**) the use of MRI compatible optical and microscope body materials, and (**3**) a custom microscope form factor to avoid camera displacement artifacts that occur during MRI gradient switching.

First, to optimize the signal quality of the BOLD fMRI images, we shielded any non-compatible electronics and materials and positioned these far from the magnetic field. For the excitation light, we coupled an external light source (488 nm laser; OBIS LX/LS Series, Coherent) into the microscope through a 400 μm optical fiber (FP400ERT, Thorlabs, **Fig 1a**). This allowed us to place the fluorescence light source outside the magnetic field of the MRI scanner. In addition, we used a shielded MRI compatible CCD camera (MRC HighResolution, MRC Systems GmbH) to record the microscope fluorescent signal. This enabled us to place the camera close to the animal’s head inside the MRI bore, which is essential for high resolution Ca^2+^ imaging.

Second, we tested all components of the microscope to avoid MRI artifacts. These included the optical lenses, microscope body, and the mouse head implant for *in vivo* imaging. **Figure 1c** shows the effect of different objective lenses, as well as the cortical window on the BOLD signal in an *ex vivo* mouse brain. We found that the magnetic susceptibility of conventional borosilicate glass (BK7) and Gradient index (GRIN) lenses for volumes typically used in microscope objectives lead to significant artifacts in the BOLD fMRI signal (for a detailed review, see 20). To resolve this issue, we designed a miniaturized microscope objective based on readily available acrylic lenses (**Fig. 1a,b**; 17-271, Edmund Optics; 36-629, Edmund Optics). This allowed us to record microscopic fluorescent images inside the MRI scanner with minimal compromise to the BOLD fMRI signal quality (**Fig. 1c, right panel**).

To further optimize the BOLD fMRI signal, we also tested different materials for the 3D printed microscope body and objective (**Supplementary Fig. 1**). We found that HDT180 (Loctite 3D 3860 HDT180, Henkel) featured high 3D printing resolution, durability, tensile strength, and resistance to heat, resulting in more durable and stable microscope recordings. In addition, HDT180 achieved SNR levels comparable to other polymers containing black pigments (**Supplementary Fig. 1h**).

Third, due to the slight magnetic receptiveness of the MRI compatible camera electronics and housing, we observed micron-sized shifts in the microscope images during the radio- frequency (RF) MR excitation pulses. To avoid such microscope motion, we measured the minimal distance (*d*) needed between the edge of the camera and the center of the magnetic field (**Fig. 1b**). We found that a distance of *d* > 6 cm was sufficient to avoid any microscope displacement (**Fig. 1d**), leading us to elongate the microscope imaging pathway by adding a two-lens relay (47-690-INK, Edmund Optics; AC080.020, Thorlabs; **Fig. 1b,d**). This effectively increased the distance between the microscope CCD camera and the RF-excitation field to 7.1 cm, resulting in stable microscope recordings (**Fig. 1b,d**).

### Simultaneous whole brain BOLD and fluorescence recordings

For *in vivo* Ca^2+^ imaging, we implanted transgenic mice (Rasgrf2-2A-dCre^21^ crossbred with Ai148D) expressing the Ca^2+^ indicator GCamp6m in cortical layer 2/3 with a round, 3 mm diameter cranial window above the barrel cortex (SSp-bfd, AP = 2.0 mm, LR = 3.0 mm).

To position and focus the MRI compatible microscope inside the MRI bore, we custom built a micrometer z-translation stage out of aluminum and 3D printed resin (VeroClear, StrataSys; **Fig. 1e**; github.com/rlemubaghs/open_mrcradle_mouse.git). We then assembled the MRI compatible microscope, the translational stage, a custom 3D printed whisker stimulation apparatus, and the housing into a cradle that fitted inside the bore of a small animal MRI scanner (**Fig. 1e**; BioSpec 70/20 USR, BRUKER BioSpin MRI GmbH).

To prepare animals for the awake imaging experiments, we started with a habituation period (>10 days) during which we handled all animals and slowly introduced them to the MRI environment and stimulation protocol (see **methods** for a detailed description). After animals became familiar with the experimental procedure and the MRI sound, we initiated the experimental phase. During imaging experiments, we individually head-fixed each mouse in the MRI cradle, mounted the MRI receiver (Rx) coil above the cortical window, positioned and focused the MRI compatible microscope and added D_2_O (D-content 99.9%; Sigma–Aldrich) as immersion media between the acrylic lens objective and the cortical window (**Fig. 1f,g**). Designing an acrylic water immersion objective was crucial as the susceptibility borders between air and tissue resulted in large signal void due to local field inhomogeneities and thus artifacts in the MR image (**Fig. 1f**, inset).

The circular field of view (FOV) of the MRI compatible microscope covered a large cortical area across the barrel cortex, and allowed us to acquire fluorescence images with single-cell resolution in the awake mouse barrel cortex inside the MRI scanner (FOV 1.45 mm^2^, pixel resolution 1.1 μm, Objective NA 0.363, optical resolution 2.5 μm +/- 0.24 μm (FWHM), **Fig. 1h,i,j**), without any noticeable BOLD fMRI artifacts or decreases in SNR.

### Simultaneous recordings of BOLD fMRI and cellular resolution calcium activity

To test and validate our combined imaging approach, we simultaneously recorded population Ca^2+^ signals and BOLD fMRI responses in awake mice during whisker stimulation (**Fig. 2**). We presented each mouse with 12 trials (block design) consisting of 10 seconds pneumonic (air) stimulation at 3 Hz (40 ms duty cycle) to the full whisker pad located contralaterally to the cortical window.

**Figure 2.**
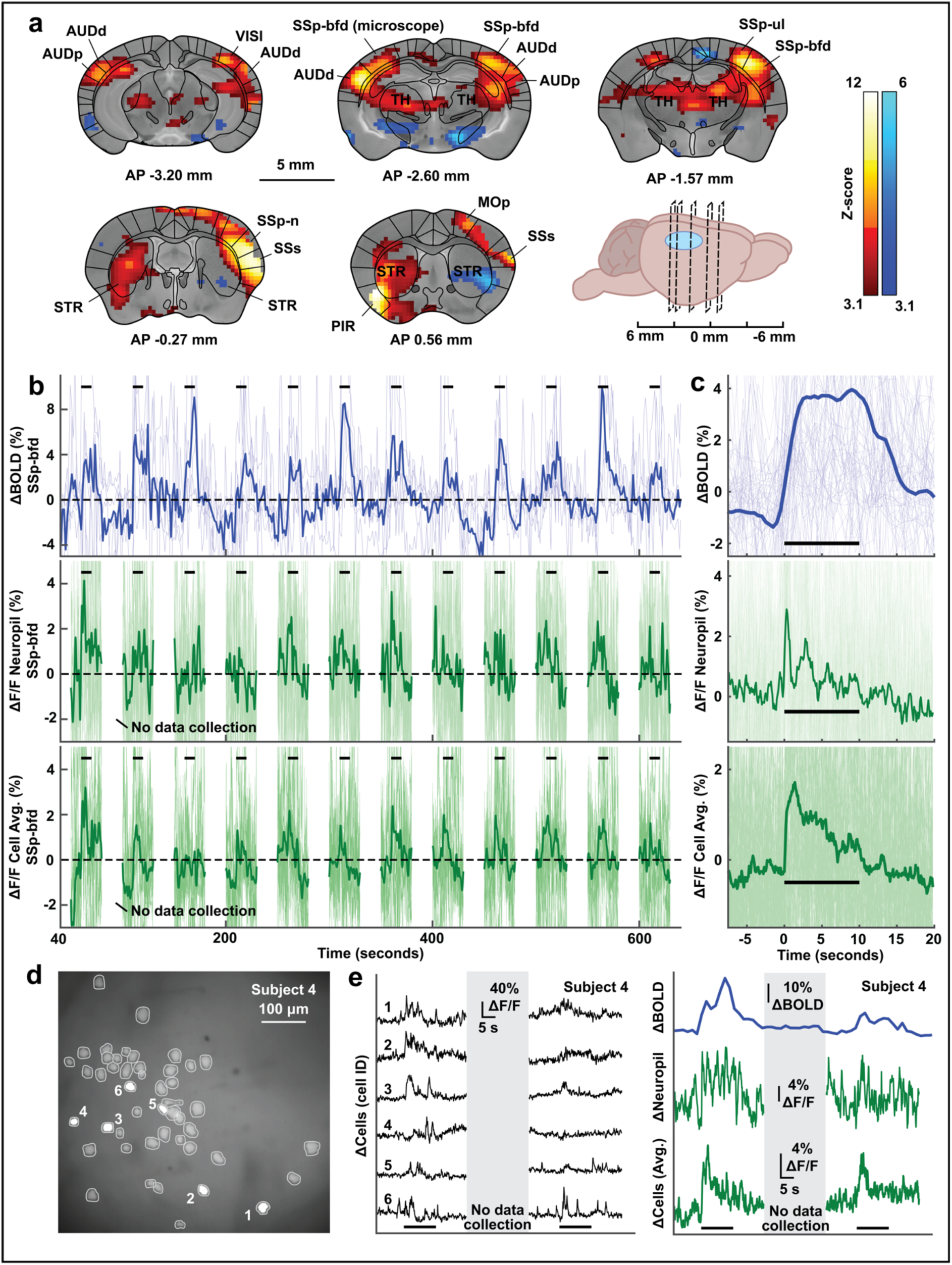
Simultaneous BOLD fMRI and calcium imaging of neural population activity at single-cell resolution. **a**, Statistical parametric map (Fixed effects, two-tailed *t*-test, z- score ≥ 3.1, FWE *P* ≤ 0.00001, N = 6) generated from the ge-EPI BOLD fMRI images acquired during contralateral whisker stimulation (N = 6 subjects). The schematic (bottom right) shows the location of each frame relative to the cortical window. Results are overlaid on the atlas described in Dorr et al., 2008 and Ioanas et al., 2021. AUDd = dorsal auditory cortex, AUDp = primary auditory cortex, MOp = primary motor cortex, PIR = piriform cortex, SSp-bfd = somatosensory barrel fields, SSp-n = somatosensory orofacial domain, SSp-ul = somatosensory upper limb domain, SSs = secondary somatosensory area, STR = striatum, TH = thalamus, VISI = primary visual cortex. **b,** BOLD fMRI (top), neuropil (middle), and cellular population Ca^2+^ activity (bottom) response in the somatosensory barrel field (SSp-bfd) averaged over animals (N = 6 subjects, 12 trials). Individual sessions are plotted in the background to show variability between subjects. Black lines indicate whisker stimulation. Interruptions in data sampling result from the laser-off period. **c,** Evoked response to contralateral whisker stimulation averaged over trials (N = 6 subjects, 12 trials). Faint lines show individual subjects. **d,** Cell map from one subject extracted with CNMF-E^26^. **e,** Example activity traces from individual cells (left), BOLD fMRI (top right), neuropil (middle right), and the neural population (bottom right) for a single mouse during whisker stimulation (black line). Black lines indicate whisker stimulation. The neural activity corresponds to the extracted cell map shown in **d**. Interruptions in data sampling result from the laser-off period.

To identify BOLD fMRI voxels that were sensitive to the stimulation we used a generalized linear model (see **methods**; **Suppl. Fig. 3**). We found robust BOLD activations across several cortical and subcortical regions in response to unilateral whisker stimulation (**Fig. 2a**; z-score ≥ 3.1, FWE *P* ≤ 0.00001, N = 6). Consistent with previous work, the main regions exhibiting BOLD signal increase included regions commonly involved in somatosensation such as the bilateral thalamus (TH), striatum (STR), secondary somatosensory area (SSs), orofacial domains (SSp-n) and the whisker domains (SSp-bfd)^22–25^. The strong activation of auditory regions, including the bilateral primary and dorsal auditory cortex (AUDp, AUDd) could be explained by the audible pneumonic pressure generated during stimulation.

To demonstrate the sensitivity of our combined imaging approach, we extracted the activity time series for all mice from significantly activated SSp-bfd voxels located ipsilateral to the microscope recordings (**Supplementary Fig. 3**; z-score ≥ 3.1, FWE *P* ≤ 0.00001, N = 6). We found that the BOLD fMRI activity showed strong stimulus-evoked potentials in the ΔBOLD signal (Wilcoxon rank-sum test *P* ≤ 0.0001; **Fig. 2b,c**). The activity also revealed a high degree of temporal signal fluctuations, most likely due to spontaneous neural activity and residual movement^12,14^. The trial-averaged response shows a well-defined hemodynamic response (∼4% signal change; Time-to-peak 80% (TTP_80%_): 2.0 seconds; **Fig. 2c**), which is in line with previous studies using a similar paradigm^12,23^.

Next, we extracted the ΔF/F Ca^2+^ activity for each individual cell (203 layer 2/3 SSp-bfd neurons; 51 ± 13 per animal; mean ± s.d.; N = 4; **Fig. 2d,e**), as well as the neuropil signal (N = 6; **Fig. 2e**) from our MRI compatible microscope recordings using CNMF-E^26^. In addition, we averaged the time series of individual cells resulting in one mean population activity trace per animal (**Fig. 2e**). For both the population and neuropil ΔF/F signal, we observed a strong stimulus-evoked signal on the subject level (Wilcoxon rank-sum test *P* ≤ 0.0001; **Fig. 2b,c**) and for the trial averaged time series (neuropil: ∼3% signal change; TTP_80%_: 0.2 seconds; population activity: ∼2% signal change; TTP_80%_: 0.8 seconds; **Fig. 2c**). Together, these results show that our new approach enables simultaneous measurements of BOLD fMRI and population activity at single-cell resolution.

### Neuronal responses exhibit distinct activation patterns based on their distance to the local vasculature

Next, we explored the potential of our new technique to investigate the local relationship between the single-cell Ca^2+^ activity and the BOLD fMRI responses. To quantify the predictive information of the SSp-bfd neural population with respect to the local BOLD fMRI signal, we used a decoding approach based on support vector machines (SVM; see **methods** for a detailed description). For this analysis, we used the ΔF/F Ca^2+^ activity traces from individual neurons to predict the average ΔBOLD fMRI signal in the SSp-bfd (**Fig. 3a**).

**Figure 3.**
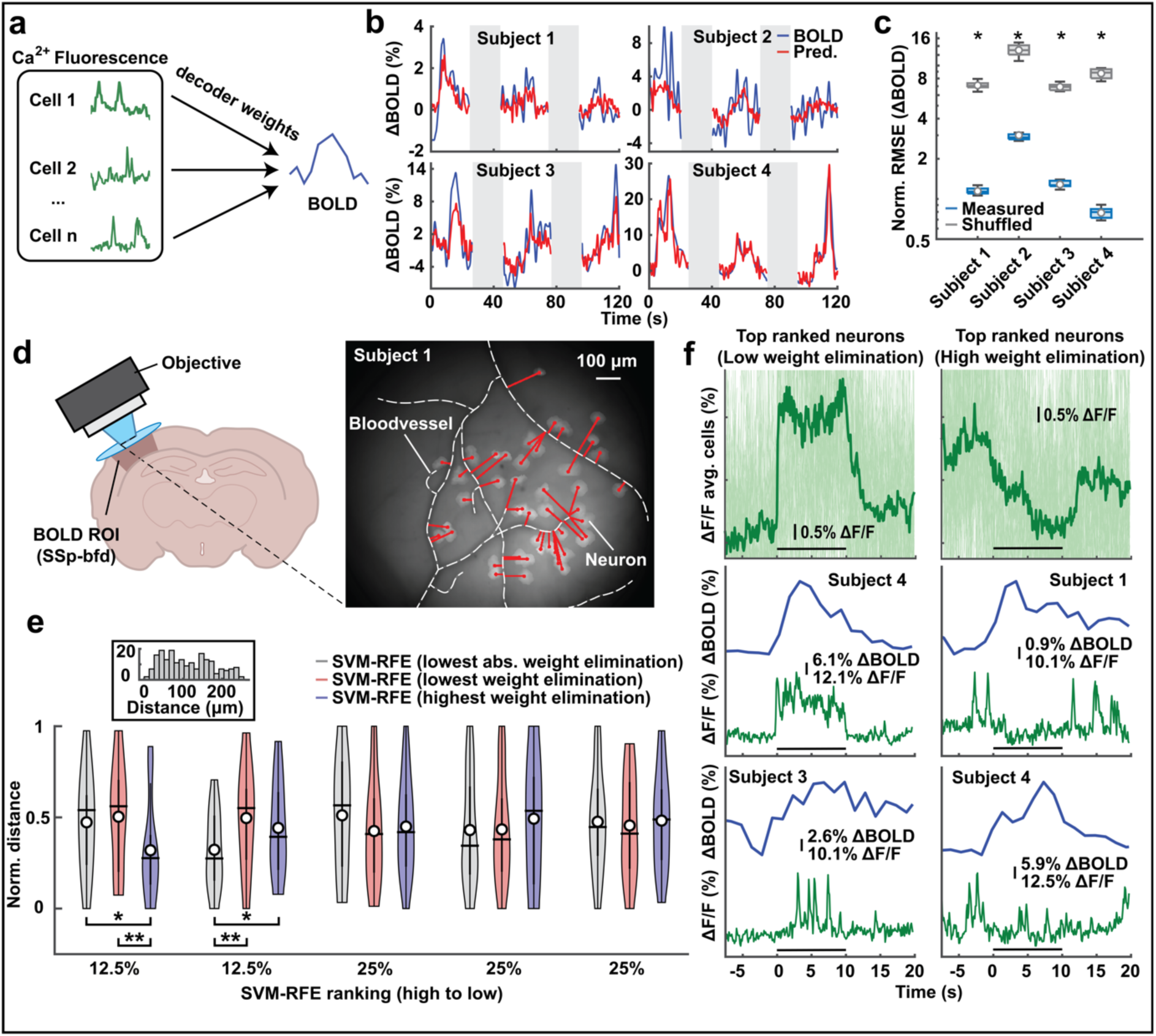
Spatial location of neurons with respect to the vascular structure determines the predictive relationship between BOLD fMRI and single-cell calcium activity. **a**, Schematic of the support vector machine decoding approach used to quantify the predictive information of the local neural population in the barrel fields (SSp-bfd) with respect to the BOLD fMRI signal. **b,** Example predictions of the decoding approach for the ΔBOLD fMRI signal in the SSp-bfd for each subject. Gray boxes indicate laser-off period. **c,** The normalized root mean square error (NMRSE) between the measured and predicted ΔBOLD fMRI signal. Boxplots show the NRMSE of the ten times cross-validated outcomes for each animal. The central line represents the median, the circle represents the mean. The minimum and maximum of the box extend to the 25th and 75th percentiles, whiskers extend to all data points. We tested the difference in performance between measured and shuffled data using a two-sample *t*-test (two-sample *t*-test; *P* ≤ 0.0001). **d,** Schematic of the experimental setting (left). We manually annotated the blood vessel patterns (right) for each animal, and defined the neuron-vascular proximity as the Euclidean 2-dimensional distance between the center of each neuron and the nearest point on the blood vessel. **e,** The neuron-vascular proximity measures, quantized based on the SVM decoding procedure in **a**. We ranked the features using recurrent feature elimination (see methods) using three individual ranking criteria (highest weights elimination, lowest weights elimination, and lowest absolute weight elimination). The central line represents the median, the circle represents the mean. The bold whisker extends to the 25th and 75th percentiles, the fine whisker extends to all data points. The width of the plot is proportional to the number of data points with the corresponding value. We tested for the difference between the distance values within each quantile using a two-sample *t*-test (two-sample *t*-test; ** = *P* ≤ 0.05; * = *P* ≤ 0.1). No correction for multiple comparisons was applied. Inset: distance histogram of all neurons. **f,** Trial average ΔF/F Ca^2+^ activity of the 12.5% highest ranked cells (top) for two ranking criteria (lowest weights elimination, left; highest weights elimination, right), and example ΔBOLD fMRI and ΔF/F single-cell Ca^2+^ activity for both ranking criteria. Black lines indicate whisker stimulation.

Our results show that the normalized root mean squared error (NRMSE) of all predicted responses is lower than the ones of the temporally shuffled baseline model (**Fig. 3b,c**; two-sample *t*-test; *P* ≤ 0.0001), and demonstrates that the neural population activity contains information about the local ΔBOLD fMRI signal.

However, population Ca^2+^ imaging provides not only information about the neural activity and vasculature structure, but also their spatial arrangement. To investigate the effect of the neurons location in predicting the local BOLD fMRI signal, we compared the Euclidean distance between the center of each neuron and the annotated vascular structure (**methods**; **Fig 3d**; **Supplementary Fig. 4**) to the contribution of each neuron to predict the BOLD fMRI signal. As before, we used SVM recurrent feature elimination (SVM-RFE; **methods**; 27,28) to rank each neuron based on their predictive information, and compared three ranking criteria: lowest absolute decoder weight elimination, lowest decoder weight elimination, and highest decoder weight elimination (see **methods** for a detailed description of the ranking criteria).

We found no difference in spatial location as compared to the whole population when neurons were ranked using the absolute predictive strength (lowest absolute decoder weight elimination; **Fig. 3e**). However, including the sign of the decoder weights in the ranking criteria revealed a difference in activity patterns between cells which was dependent on their location with respect to the vasculature (**Fig. 3e,f**; **Supplementary Fig. 5**). We found that neurons with a strong response to contralateral whisker stimulation were exclusively assigned positive decoder weights (lowest decoder weight elimination; **Fig. 3f**; left), and showed a heterogeneous pattern in their distance from the vasculature that did not differ from the overall distribution (**Fig. 3e**; two-sample *t*-test; *P* = 0.363). In contrast, neurons that were assigned negative decoder weights (highest decoder weight elimination) showed a decreased response during whisker stimulation (**Fig. 3f**; right) and were located significantly closer to the vascular structure (**Fig. 3e**; two-sample *t*-test; *P* = 0.015).

These results show that subpopulations of neurons in layer 2/3 of the SSp-bfd contain distinct information about the vascular BOLD fMRI response. In addition, we leveraged the spatial component of the MRI compatible microscope recordings to show that the direction of the neuronal response was dependent on the distance to the local blood vessel structures. In the following section we will present a second experimental approach of how our new technique enables researchers to investigate the properties of local circuit computation within the context of global brain activity.

### From local population measurements to large-scale recordings of brain connectomes

Local neuronal activity (e.g., in the SSp-bfd) is influenced by inputs from multiple brain regions through direct and indirect afferent connections^29,30^. To fully understand local computation, it is therefore critical to simultaneously monitor activity in connected brain areas that are often distributed throughout the brain. In this section, we demonstrate how our combined Ca2+ fMRI BOLD imaging approach can be leveraged to examine the predictive information of the hemodynamic BOLD fMRI activity from different regions with respect to local population Ca^2+^ activity in the SSp-bfd.

As before, we utilize a decoding approach based on SVM (see **methods**). However, in contrast with the previous analysis we used the significantly activated voxels in the SSp-bfd (z-score ≥ 3.1, FWE *P* ≤ 0.00001) to predict the average ΔF/F neural population activity low-pass filtered at 0.01 Hz (**Fig. 4a**). To avoid overfitting due to the high number of significantly activated voxels in the SSp-bfd, we applied PCA to the BOLD fMRI data to decrease the number of input dimensions (13 components explained ≥ 90% of the variance) into the decoder.

**Figure 4.**
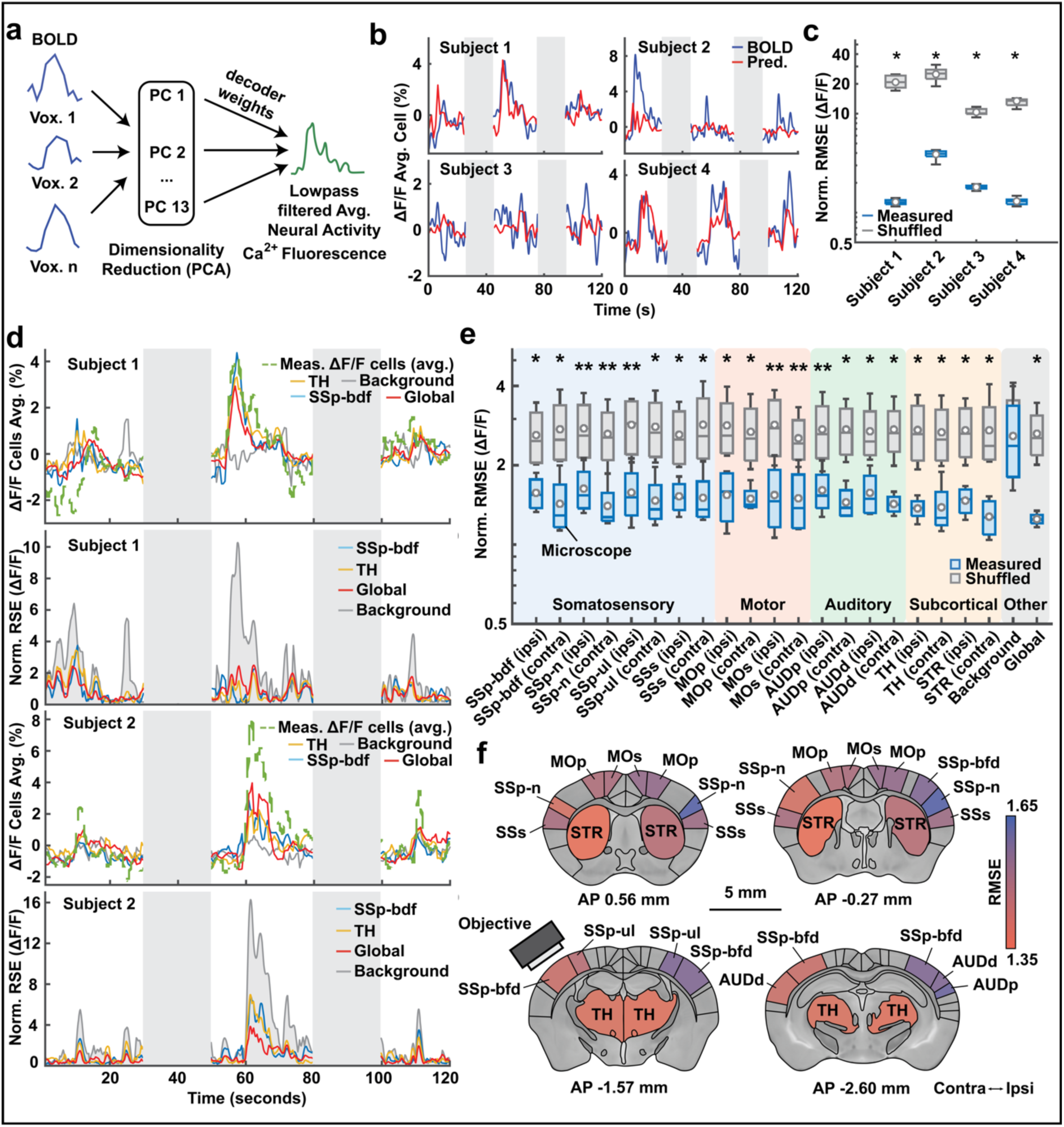
Predictive power of BOLD fMRI on the SSp-bfd population activity varies across different brain regions involved in whisker stimulation. **a**, Schematic of the SVM decoding approach to predict the BOLD fMRI signal of the barrel fields (SSp-bfd) from the local SSp-bfd Ca^2+^ population activity. **b,** Example trial predictions of the BOLD signal from the Ca^2+^ population activity in the SSp-bfd. Gray boxes indicate laser-off period. **c,** Normalized root mean square error (NMRSE) between the measured and predicted ΔF/F population activity. Boxplots show the NRMSE of the ten times cross-validated outcomes for each animal. The central line represents the median, the circle represents the mean. The minimum and maximum of the box extend to the 25th and 75th percentiles, whiskers extend to all data points. We tested for the difference between the measured and shuffled data using a two-sample *t*-test (two-sample *t*-test; * = *P* ≤ 0.0001). **d,** Example predictions of the SSp-bfd ΔF/F population activity (upper) and corresponding normalized root square error (below; NRSE) for different brain areas and background noise. Gray boxes indicate laser-off period. SSp-bfd = somatosensory barrel fields, TH = thalamus. **e,** NMRSE between the ΔF/F and predicted ΔF/F population activity for areas involved in whisker stimulation. Boxplots show the NRMSE from all animals. The central line represents the median, the circle represents the mean. The minimum and maximum of the box extend to the 25th and 75th percentiles, whiskers extend to all data points. We tested the difference in performance between measured and shuffled data using a Wilcoxon rank-sum test (Wilcoxon rank-sum test; * = *P* ≤ 0.1; ** = *P* ≤ 0.05). AUDd = dorsal auditory cortex, AUDp = primary auditory cortex, MOp = primary motor cortex, MOs = secondary motor cortex, SSp-bfd = somatosensory barrel fields, SSp-n = somatosensory orofacial domain, SSp-ul = somatosensory upper limb domain, SSs = secondary somatosensory area, STR = striatum, TH = thalamus. Contralateral and ipsilateral indicated with respect to the whisker stimulation. **f,** Coronal magnitude map of the mouse brain illustrating the predictive information of the ΔBOLD in relation to the neural population activity based on the NRMSE described in **e**.

Our results show that the NRMSE of all predicted responses fall far below the temporally shuffled baseline model (**Fig. 4b,c**; two-sample *t*-test; *P* ≤ 0.0001). This demonstrates that the ΔBOLD fMRI signal contains information about the local neural population activity.

Next, we extended this approach to different brain regions previously implicated in whisker stimulation^29,31^ (**Fig. 2a**). To avoid overestimating the amount of information for larger anatomical regions due to overfitting of the SVM, we randomly subsampled 50 voxels from each region before applying PCA. To cross-validate our results, we repeated this procedure ten times for each animal, and averaged the decoder performance to produce one value per region^32,33^. To ensure that vascular information from preceding or delayed regions was represented within the training set, we also included bidirectional time-delayed ΔBOLD activity time points (-1, 0.5, 0, 0.5, and 1 second).

Our results show that the local neural population activity in the SSp-bfd can be predicted better from the ΔBOLD fMRI signal across cortical and subcortical regions involved in whisker stimulation^29,31^ as compared to the temporally shuffled baseline model (**Fig. 4d,e**; Wilcoxon rank-sum test; * = *P* ≤ 0.1; ** = *P* ≤ 0.05; **Supplementary Fig. 6**). When we visualized the performance of each region through a magnitude map (**Fig. 4f**), we observed a significant difference in predictive power between regions contralateral to the stimulation as compared to ipsilateral (two-sample *t*-test; *P* = 0.034). However, we found no significant differences for individual regions.

Together these results demonstrate the new possibilities of simultaneous global BOLD fMRI and cellular-resolution Ca^2+^ imaging. We show that different cortical and subcortical regions contain relevant predictive information about neural activity in the SSp-bfd. Interestingly, these regions are consistent with previous accounts of the mouse whisker system^22–25^, and the anatomical connections between these regions^29,31^.

## Discussion

In this work, we developed an MRI compatible microscope that enables the simultaneous imaging of single-cell Ca^2+^ activity during brain-wide BOLD fMRI acquisition. Previously, researchers combined optical imaging methods, such as fiber photometry-based Ca^2+^ imaging, and BOLD fMRI^15–17,19^. However, due to the limited spatial resolution of fiber-based methods, (non-)neuronal cellular and neuropil signals are grouped together. Here, we demonstrate that the resolution of our MRI compatible microscope is capable of measuring the calcium signals of a population of single cells within the MRI scanner. Similarly, combined electrophysiology and BOLD fMRI allows measuring neural signals with single-cell resolution during BOLD fMRI acquisition, and several groups have used this method to investigate the neural underpinnings of the BOLD fMRI signal^34^. However, compensating for the electromagnetic interference inside the MRI scanner is costly and technically very demanding. In addition, several key questions such as cell specific interactions, for example, between neuronal and glial cells and the BOLD fMRI signal, are difficult to address without the genetic targeting that optical imaging methods offer^35^. Our approach allows for genetic targeting, and in addition is cost-efficient and straightforward to implement as the microscope is easy to build and to install inside the MRI scanner.

Nevertheless, designing a fMRI compatible microscope brings additional challenges. To overcome these, we (**1**) designed the MRI compatible optics and tailored microscope form factor, (**2**) tested and exclusively used MRI compatible materials for uncomplicated manufacturing, and (**3**) appropriately shielded or positioned all necessary MRI incompatible microscope electronics. In a first proof of principle experiment, we demonstrated that our approach results in a high signal-to-noise ratio for BOLD fMRI images without compromising fluorescent signal transmission. Next, we demonstrated the potential of combined BOLD fMRI and Ca^2+^ activity population imaging in two proof-of-concept experiments in awake animals by investigating (**1**) how the spatial relationship between local SSp-bfd neurons and the vasculature relates to the local BOLD fMRI signal, and (**2**) how the activity from regions across the brain relates to the local neural activity in the SSp-bfd.

In the first application example, we used the simultaneous local BOLD fMRI and single-cell calcium activity in the SSp-bfd to investigate the effect of neuron location in predicting the local BOLD fMRI signal. We used SVM-RFE to rank each neuron based on its predictive performance and found a difference in activity patterns between cells which was dependent on their location with respect to the vasculature. We observed that highly predictive neurons closer to the vasculature exhibited a negative neural response during whisker stimulation, contrasting with the positive vascular BOLD fMRI activity. One potential explanation of the negative activity is that it is related to their connectivity with vasoactive interneuron subtypes. Previous studies suggest that inhibitory neurons are involved in neurovascular coupling^36–40^, through the release of neurotransmitters that exert strong vasoactive effects when released close to the vasculature (vasodilatory VIP^+^: 41; vasoconstricting SOM^+^: 41,42). It could therefore be possible that excitatory neurons which exhibit a negative relationship with respect to the local BOLD fMRI activity are embedded in a circuit that integrates these vasoactive interneurons. Using our approach, future studies can now investigate the relationship between the hemodynamic response and specific inhibitory subclasses at the single-cell level in more detail by leveraging modern genetic targeting methods. Moreover, our microscope can be easily adapted to enable the simultaneous manipulation and recording of two genetically distinct cell populations utilizing multicolor imaging^43,44^, or the integration of optogenetic stimulation during Ca^2+^ fluorescence imaging^45,46^.

In the second application example, we investigated the influence of direct and indirect afferent connections of different brain regions on the local calcium activity of the neural population in the SSp-bfd. We found that the neural population activity in the SSp-bfd could be predicted with a high accuracy from the BOLD fMRI activity in different regions of the mouse brain. This finding is consistent with previous studies in that regions implicated in whisker stimulation^22–25^, as well as anatomically connected areas^29,31^ contained a high level of information about the local neural activity in the SSp-bfd. Using our method, future studies can now investigate whether regions across the brain contain distinct predictive information about diverse local neuronal subsets that differ in their genetics or projection specificity. In addition, the suitability of our microscope to enable awake rodent fMRI experiments means that these studies can be conducted during more natural behavioral paradigms.

Ultimately, a better understanding of the relationship between neurons and the vasculature will lead to a better interpretability of animal and human BOLD fMRI studies^47–49^, especially when combined imaging experiments are performed in awake animals^23,50–52^. In addition, combined local and global activity measurements will enable a better understanding of the functional organizing principles of the brain across scales.

## Methods

All procedures were authorized by the veterinary office of the Canton Zurich, Switzerland, and are in agreement with the guidelines published in the European Communities Council Directive of November 24, 1986 (86/609/EEC).

### Animals

Mice were kept on a 12/12-hour light cycle with unlimited access to food and water. In total, 8 male mice aged 4 - 6 months (25-35 g) were used for experiments. We collected all data from Rasgrf2-2A-dCre mice (21; Jackson Laboratory, stock number 022864) crossbred with Ai148D mice (Jackson Laboratory, stock number 030328). The resulting mice expressed GCaMP6m in cortical layer 2/3 pyramidal neurons (**Supplementary** Fig. 2).

### Surgical Procedures

During all steps of the surgical procedure, we ensured MR-compatibility of all materials, as well as the health of the tissue in order to optimize the SNR for both imaging modalities.

#### Anesthesia

For all procedures, we administered buprenorphine (Bupaq, Streuli; 0.1 mg/Kg) to all animals 30 minutes prior to anesthesia. We induced anesthesia with 5% Isoflurane and maintained it using 1.5-2% during surgery. After we head-fixed the mouse in the stereotaxic frame (Kopf Instruments), we removed the hair around the planned incision using dilapidation cream. During the whole surgical procedure, mice received 95% medical grade O2 (Pangas, Conoxia) through a face mask, while we maintained the body temperature at 37 degrees Celsius using a temperature controller and a heating pad.

#### Cortical implant

Before opening the cranium, we injected 2% Lidocaine between the skin and the skull as local anesthetic (Lidocaine, Streuli Pharma AG). After removal of the skin, we cleared the skull surface of soft tissue and debris. To increase the stability and attachment of the implant, we then applied a thin layer of phosphoric acid gel (etching gel; DMG Dental-Material GmbH) and removed it after 30 seconds with Phosphate Buffered Saline (PBS, Sigma–Aldrich). In addition, we removed excess blood and applied a thin layer of Scotchbond universal adhesive (Scotchbond, 3M) to stabilize the exposed skull during the craniotomy and to create a barrier between the skull and the skin.

Finally, to gain optical access to the cortex, we made a 3 mm diameter craniotomy above the region of interest (SSp-bf; AP = 2.0 mm, LR = 3.0 mm), and sealed it with a single borosilicate glass cranial window (3 mm diameter; **Fig. 1f**). After fixing the cortical window in place using UV and moisture-curable adhesive (Loctite 4305, Henkel), we placed a custom head-bar (Polyether ether ketone, PEEK; **Fig. 1f**) around the window and attached it with Scotchbond.

#### Analgesic regime

For 3 days after each surgical procedure animals received Buprenorphine s.c. (Bupaq; Streuli, 0.1 mg/Kg) every 6 h during the light day cycle and in the drinking water (Bupaq; Streuli, 0.01mg/mL) during the dark day cycle as well as Carprofen s.c. (Rimadyl; Zoetis, 4 mg/Kg) every 12 h.

### Habituation procedure for combined imaging in awake animals

Due to the slow acquisition of BOLD fMRI, movement can cause signal displacement within an acquisition cycle. In addition, neuromodulatory factors such as stress have been shown to have a strong effect on neurovascular coupling, leading to confounding effects in the BOLD signal^53,54^. To minimize possible confounding factors, we implemented a rigorous habituation protocol.

After the surgery, we started to handle animals twice a day for a period of at least 3 minutes. Once animals were fully recovered, we initiated the habituation period. The duration of the habituation period depended on the behavioral performance of each individual animal, and lasted a minimum of 10 days. During the first week, we then started to restrain animals in a mock MRI machine once a day for two to three days followed by one day of rest. We set the duration of the head fixation to start at 10 minutes, and stepwise increased it by 5 minutes until the average duration of a full experiment was reached (30 minutes). We introduced MRI scanner sounds during the fourth day of habituation, and gradually increased over 4 sessions till it reached the same pressure level as experienced during an MRI session (∼95 dB). In addition, we used a blue ambient light to mimic the on-off cycles of the MR-compatible microscope excitation light.

Once the animal showed no discernible response to the head fixation, sound pressure level, and excitation light^55^, we continued the habituation inside the MRI machine. The habituation sessions consisted of a functional scan that included the full stimulation protocol and a structural scan. We used the framewise displacement (FWD)^56^, to evaluate habituation, and we considered it as successful when the average FWD for a full session was below 0.05 mm, and at least 50% of trials showed no excessive motion (FWD spikes > 0.075 mm). We chose the values for the FWD thresholds based on the BOLD fMRI acquisition sequence, where FWD spikes should not exceed more than 50% of the voxel size due to spill over of the signal to nearby voxels. Once habituation was considered successful (> 3 continuous days of successful sessions), we initiated the combined imaging experiments. We did not initiate data collection in two animals because they did not reach the specified criteria.

### Assembly and components of the MRI compatible microscope

We assembled all optics using a custom 3D printed housing (Loctite 3D 3860 HDT180, Henkel; Objet 30 Pro, Stratasys; **Fig. 1a**) and fixed them in place using low-staining optical glue (NOA61, Norland). We used durable and high temperature resistant resin to avoid warping due to pressure and the surface heating of the camera. We found HDT180 to be the optimal tradeoff between printer resolution, durability, tensile strength and heat-resistance (**Supplementary** Fig. 1).

In the excitation pathway, blue light (wavelength λ = 488nm) originating from the tip of a multimode fiber coupled to an off-site laser (OBIS LX/LS Series, Coherent; **Fig. 1b**) is collimated using a 5 mm double convex lens (63-594-ink, Edmund Optics), and passes through a diffuser with a scatter angle of 10 degrees (edc-10, Rochester). The collimated light is combined with the optical path through a dichroic mirror (ET Dualband beamsplitter FITC/CY3, AHF) before it is focused on the brain surface through the objective.

In the imaging pathway, the emitted fluorescence is collected by the custom objective consisting of three small diameter acrylic aspheric lenses (17-271, Edmund Optics; 36-629, Edmund Optics). In order to minimize the vertical footprint of the microscope, the optical path is diverted 90 degrees by a mirror (350-700 nm imaging, AHF). Along the horizontal axis, light is relayed by two achromatic lenses (47-690-INK, Edmund Optics; AC080.020, Thorlabs). Finally, a second mirror reflects the image downwards through an emission filter (ET Dualband Emitter FITC/CY3, AHF) onto the MRI compatible camera (MRC HighResolution, MRC Systems GmbH; 1280x960 pixels at 5.3 μm pitch).

### Imaging parameters

#### Functional and structural MRI parameters

We acquired MRI images on a 7T/11 cm horizontal bore scanner (BioSpec 70/20 USR, BRUKER BioSpin MRI GmbH) with a BGA 9SHP (90mm, max 750 mT/m gradient strength, slew rate 6840 T/m/s, BRUKER BioSpin MRI GmbH). The scanner was fitted with a RF RES 300 1H 089/072 QSN TR AD volume coil combined with an RF SUC 300 1H LNA AV3 surface coil to receive the radiofrequency signal (BRUKER BioSpin MRI GmbH). BOLD fMRI data was acquired using a single-shot gradient-echo (GE) echo planar imaging (EPI) pulse sequence (TE/TR: 15/1500, BW: 250 kHz, FA: 60, FOV: 20/10, Image Size: 100/50, In Plane resolution: 0.2/0.2/0.8 mm, Slices: 7, Slice Gap: 0.1). In addition, a T2-weighted 3D TurboRARE (TE/TR: 21.86/2500, FOV: 20/10, Image Size: 180/120, we acquired the in-plane resolution: 0.111/0.083/0.8 mm, Slices: 7, Slice Gap: 0.1) for image registration.

#### Calcium imaging parameters

During BOLD fMRI we simultaneously measured Ca^2+^ activity of a cell population within the barrel cortex. We acquired frames of 1280x960 pixels at 12 bit with a frame rate of 10 Hz using custom C++ code. To acquire the Ca^2+^ imaging data we used laser intensities between 0.1-0.3 mW depending on the strength of the GCaMP6f expression. We streamed all recorded data directly to the hard-drive of a desktop computer and synchronized with the recorded stimulus times.

## Data processing

### Functional and structural MRI data preprocessing

We collected and stored all MRI data in the proprietary Bruker ParaVision format, and consequently converted to BIDS through the Bruker-to-BIDS repositioning pipeline of the SAMRI package^57^. After we converted the data to the BIDS format, the Generic registration workflow was used to preprocess the data^57,58^. Within this pipeline, we preprocessed the data by performing motion and slice timing corrections^59^, bias field correction^60^, registration to an atlas^61^, and spatiotemporal smoothening (spatial smoothing in coronal plane up to 200 µm; low-pass filtered in the temporal domain with a period threshold of 100 seconds).

### Generalized linear model and time series extraction for BOLD fMRI data

We modeled volumetric data using FSL (62; version 6.0.4), through subject level regression. We used six motion parameters and their first derivatives as covariates in the model. We used subject level contrasts and variance estimates as input into the mixed effects group level analysis using a gaussian hemodynamic response function (phase = 0s, sigma = 2.8s, peak lag = 5s). We extracted the average time series over all significantly activated voxels after cluster correction (z-score ≥ 3.1, FWE *P* ≤ 0.00001) and re-expressed in units of relative change, given by ΔBOLD(t) = (BOLD(t) - BOLD_0_)/BOLD_0_, where BOLD_0_ is the median filtered value (window size of 180 seconds) at each timestep. Due to the difference in temporal sampling rate between Ca^2+^ imaging and BOLD fMRI, we used linear interpolation to increase the data points of the ΔBOLD signal to match the 100 ms acquisition interval of the ΔF/F Ca^2+^ activity.

### Calcium imaging preprocessing

We implemented the following procedure to preprocess the video of each individual animal. First, we manually determined a region of interest which contained fluorescence, and spatially downsampled all frames by a factor of 2. Next, frames that did not contain a fluorescent signal as a result of excitation laser deactivation between stimulus presentations were removed, and TurboReg image registration algorithm^63^ was used to account for spatial translations by aligning each frame to a reference frame. Finally, we temporally downsampled the video by a factor of 2, resulting in a frame rate of 5 Hz.

#### Cell extraction and validation

We extracted cellular activity and neuropil traces using the CNMF-E pipeline^26^, using the following parameters: gSig = [5,5], gSiz = [17,17], ring_radius = 2, min_corr = 0.85, min_pnr = 8, deconvolution = constrained FOOPSI algorithm, decay_time = 0.4. We evaluated the spatial and temporal components of every extracted unit based on neuron shape, stability of the neural trace, temporal evolution of identified events (Ca^2+^ indicator dynamics), and the consistency of the identified events over the session, and outliers (obvious deviations from the normal distribution) were discarded. We were able to extract a total of 203 layer 2/3 SSp-bfd neurons (51 ± 13 per animal; mean ± s.d.; N = 4) from the MRI compatible microscope recordings. We were unable to extract cells from two animals leading us to exclude them from analysis that involved single cell Ca^2+^ activity.

#### Annotation of the vascular structure and neuron-vascular proximity

To estimate the proximity of neurons to the vasculature, we manually annotated the blood vessel patterns for each animal (**Supplementary** Fig. 4). We used the time averaged and band pass filter (100-150 Hz) microscopic field to gain clear visibility of the vascular structure. Special care was taken to exclude features that were not part of the blood vessel formation. All annotations were validated by two additional researchers.

Next, we estimated the Euclidean distance between the center of each cell and the vascular structure via:

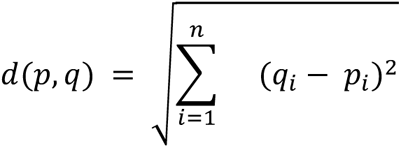

where p and q are two points in Euclidean n-space. Note that this description of the Euclidean distance is restricted to a two-dimensional field due to the limited capacity of single-photon microscopy to resolve depth. Finally, we used the minimal distance value for each individual neuron to determine the neuron-vascular proximity metric.

#### Support vector machines (SVM) and recurrent feature elimination (SVM-RFE)

To quantify the predictive information of both imaging modalities with respect to the other modality, we used a decoding approach based on linear epsilon-insensitive SVM regression models. For each model, the decoding performance was quantified by the Root Mean Square Error (RMSE) according to:

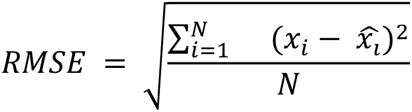

where x_i_ is the observed ΔBOLD fMRI time series and 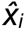 the estimated ΔBOLD fMRI time series. Subsequently, the RMSE was normalized by the standard deviation (*σ_x_*) to compare error levels between animals:

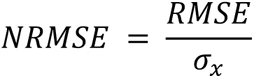

To estimate the baseline model, we repeated this procedure with temporally permuted training features, providing an upper bound on the possible NRMSE given the prediction features.

In addition, we used recurrent feature elimination to rank the predictive features based on their level of contribution in predicting the alternate imaging modality. SVM-RFE uses the weight magnitude as a ranking criterion to remove less important features recursively^27,28^, providing a subset of features that achieve the highest performance. First, an SVM regression model is trained using all available features. The resulting *β*-weights are then ranked based on a predefined criterion. The feature with the lowest ranking is eliminated, and given the lowest remaining rank. The process is then repeated without the eliminated feature until all features have received a ranking.

In the current paper we used three ranking criteria yielding three distinct rankings. 1) Lowest absolute decoder weight elimination. Weights are ranked based on the absolute *β*-weights, and the lowest value is iteratively removed. This procedure does not take into account the sign of the assigned *β*-weights, providing both positive and negative *β*-weights an equal chance at receiving a high ranking. 2) Lowest decoder weight elimination. Weights are ranked based on the magnitude of the *β*-weights, and the lowest value is iteratively removed. This procedure results in a higher chance for *β*-weights with a positive value to receive a high ranking. 3) Highest decoder weight elimination. Weights are ranked based on the magnitude of the *β*-weights, and the highest value is iteratively removed. This procedure results in a higher chance for *β*-weights with a negative value to receive a high ranking.

### Software and Statistical Tests

For all data analysis (image pre-processing and population analysis) and statistics we used custom MATLAB and Python code. The sample sizes required for this study were initially estimated based on pilot behavior studies, but we did not use formal statistical tests to predetermine sample size. We excluded one animal because we observed no neuronal activity and low SNR of the MRI images, possibly due to degradation of the implant over time. We excluded one animal due to significant air pockets trapped in the glue close to the brain.

### Validation of imaging methodology

#### Perfusion

After completion of the experiments, we administered a terminal anesthesia with Pentobarbital (Esconarkon; Streuli, 200 mg/Kg) and perfused transcardially with PBS followed by 4% paraformaldehyde (PFA). We removed brain tissue and post-fixed it for 24-48 h in 4% PFA. We then prepared coronal slices (50 μm thick) on a Vibratome (VT1000 S; Leica) and stored these in PBS.

#### Verification of cell type

We used standard immunofluorescence protocols to stain inhibitory and excitatory neurons. We incubated slides overnight with either the primary antibody rabbit anti-Neurogranin (1:2000, 07-425 Millipore) or rabbit anti-GAD65 (1:500, AB1511, Millipore) at 4°C, followed by a 2-hour incubation at room temperature with the secondary antibody Alexa 594 anti-rabbit (1:200, A-11062 Invitorgen). We further stained slides for 4 min with DAPI (1:1000, D1306, Invitrogen) in PB (0.1M) prior to mounting. We then acquired confocal fluorescence images in the red (594 nm, Neurogranin or GAD65), green (488 nm, GCaMP6m), and blue channel (390 nm, DAPI). Finally, we evaluated the resulting images with respect to their labeling overlap (see **Supplementary** Fig. 2).

### Data Availability Statement

Data used to reproduce figures will be provided with this paper and made publicly available via Figshare. All other data (raw imaging data and pre-processed activity data) will be made available upon request.

### Code Availability Statement

All (optical) design files and code related to the experimental setup (microscope and behavioral setup) will be made publicly available upon publication. The MATLAB and Python code scripts detailing all aspects of the performed analysis will be made available upon request.

## Acknowledgements

We would like to thank Harald Dermutz for fruitful discussions and for feedback on the design of the study and the interpretation of the results. We would like to thank Simone Holler for the preparation of the histology slides. Finally, we would like to thank Silvio Scherr, Lukas Lüchinger and Fabian Eggimann for their help with the manufacturing of the microscope and behavioral apparatus. Valerio Zerbi is supported by the Swiss National Science Foundation (SNSF) ECCELLENZA fellowship (PCEFP3_203005). This work is supported by the Swiss National Science Foundation (B.F.G. CRSII5-173721 and 315230 189251), ETH project funding (B.F.G. ETH-20 19-01) and the Human Frontiers Science Program (RGY0072/2019).

## Author Contributions

R.U. and B.F.G. conceptualized the study, R.U. carried out the animal preparation, in vivo imaging experiments, setup and microscope design, and data analysis, R.B. helped with animal preparation, in vivo experiments, and data analysis, H.K.H helped with the lens design, M.M. helped with the MRI pilot experiments and data analysis, F.Y. and V.Z. provided conceptual feedback for the behavior experiments, data acquisition and data analysis, R.U. and B.F.G wrote the manuscript.

## References

1. Uludağ, K., Müller-Bierl, B. & Uğurbil, K. An integrative model for neuronal activity-induced signal changes for gradient and spin echo functional imaging. Neuroimage 48, (2009).

2. Buxton, R. B. Interpreting oxygenation-based neuroimaging signals: the importance and the challenge of understanding brain oxygen metabolism. Front. Neuroenergetics 2, (2010).

3. Polimeni, J. R. & Lewis, L. D. Imaging faster neural dynamics with fast fMRI: A need for updated models of the hemodynamic response. Prog. Neurobiol. 207, 102174 (2021).

4. Gong, Y. et al. High-speed recording of neural spikes in awake mice and flies with a fluorescent voltage sensor. Science 350, 1361–1366 Preprint at 10.1126/science.aab0810 (2015).

5. Cardin, J. A., Crair, M. C. & Higley, M. J. Mesoscopic Imaging: Shining a Wide Light on Large-Scale Neural Dynamics. Neuron 108, 33–43 (2020).

6. de Vries, S. E. J. et al. A large-scale standardized physiological survey reveals functional organization of the mouse visual cortex. Nat. Neurosci. 23, (2020).

7. Demas, J. et al. High-speed, cortex-wide volumetric recording of neuroactivity at cellular resolution using light beads microscopy. Nat. Methods 18, 1103 (2021).

8. Cramer, S. W. et al. Through the looking glass: a review of cranial window technology for optical access to the brain. J. Neurosci. Methods 354, 109100 (2021).

9. Goldey, G. J. et al. Removable cranial windows for long-term imaging in awake mice. Nat. Protoc. 9, (2014).

10. Kim, T. H. & Schnitzer, M. J. Fluorescence imaging of large-scale neural ensemble dynamics. Cell 185, (2022).

11. Wapler, M. C. et al. MR-compatible optical microscope for in-situ dual-mode MR- optical microscopy. PLoS One 16, e0250903 (2021).

12. Desjardins, M., et al. Awake Mouse Imaging: From Two-Photon Microscopy to Blood Oxygen Level–Dependent Functional Magnetic Resonance Imaging. Biological Psychiatry: Cognitive Neuroscience and Neuroimaging 4, 533–542 (2019).

13. Scaglione, A., Moxon, K. A., Aguilar, J. & Foffani, G. Trial-to-trial variability in the responses of neurons carries information about stimulus location in the rat whisker thalamus. Proc. Natl. Acad. Sci. U. S. A. 108, (2011).

14. Mateo, C., Knutsen, P. M., Tsai, P. S., Shih, A. Y. & Kleinfeld, D. Entrainment of Arteriole Vasomotor Fluctuations by Neural Activity Is a Basis of Blood-Oxygenation-Level-Dependent ‘Resting-State’ Connectivity. Neuron 96, (2017).

15. Schlegel, F. et al. Fiber-optic implant for simultaneous fluorescence-based calcium recordings and BOLD fMRI in mice. Nat. Protoc. 13, 840–855 (2018).

16. Schulz, K., et al. Simultaneous BOLD fMRI and fiber-optic calcium recording in rat neocortex. *Nature Methods* vol. 9 597–602 Preprint at 10.1038/nmeth.2013 (2012).

17. Liang, Z., Ma, Y., Watson, G. D. R. & Zhang, N. Simultaneous GCaMP6-based fiber photometry and fMRI in rats. J. Neurosci. Methods 289, 31–38 (2017).

18. Schwalm, M. et al. Cortex-wide BOLD fMRI activity reflects locally-recorded slow oscillation-associated calcium waves. Elife 6, (2017).

19. Lake, E. M. R. et al. Simultaneous cortex-wide fluorescence Ca2+ imaging and whole-brain fMRI. Nat. Methods 17, 1262–1271 (2020).

20. Wapler, M. C. et al. Magnetic properties of materials for MR engineering, micro-MR and beyond. J. Magn. Reson. 242, (2014).

21. Harris, J. A., et al. Anatomical characterization of Cre driver mice for neural circuit mapping and manipulation. *Front. Neural Circuits* 8, (2014).

22. Esmaeili, V. et al. Cortical circuits for transforming whisker sensation into goal-directed licking. Curr. Opin. Neurobiol. 65, (2020).

23. Chen, X. et al. Sensory evoked fMRI paradigms in awake mice. Neuroimage 204, (2020).

24. Reig, R. & Silberberg, G. Multisensory Integration in the Mouse Striatum. Neuron 83, 1200 (2014).

25. Zareian, B., Lam, A. & Zagha, E. Dorsolateral Striatum is a Bottleneck for Responding to Task-Relevant Stimuli in a Learned Whisker Detection Task in Mice. J. Neurosci. 43, (2023).

26. Zhou, P. et al. Efficient and accurate extraction of in vivo calcium signals from microendoscopic video data. eLife 7:e28728. (2018).

27. Guyon, I., Weston, J., Barnhill, S. & Vapnik, V. Gene Selection for Cancer Classification using Support Vector Machines. Mach. Learn. 46, 389–422 (2002).

28. Cho, B. H. et al. Application of irregular and unbalanced data to predict diabetic nephropathy using visualization and feature selection methods. Artif. Intell. Med. 42, (2008).

29. Aronoff, R. et al. Long-range connectivity of mouse primary somatosensory barrel cortex. Eur. J. Neurosci. 31, (2010).

30. Feldmeyer, D. Excitatory neuronal connectivity in the barrel cortex. Front. Neuroanat. 6, (2012).

31. Yamashita, T. et al. Diverse Long-Range Axonal Projections of Excitatory Layer 2/3 Neurons in Mouse Barrel Cortex. Front. Neuroanat. 12, (2018).

32. Zhong, Y. et al. Detecting functional connectivity in fMRI using PCA and regression analysis. Brain Topogr. 22, (2009).

33. Sobczak, F., Pais-Roldán, P., Takahashi, K. & Yu, X. Decoding the brain state-dependent relationship between pupil dynamics and resting state fMRI signal fluctuation. Elife 10, (2021).

34. Logothetis, N. K., Pauls, J., Augath, M., Trinath, T. & Oeltermann, A. Neurophysiological investigation of the basis of the fMRI signal. Nature 412, (2001).

35. Attwell, D. et al. Glial and neuronal control of brain blood flow. Nature 468, 232–243 (2010).

36. Krawchuk, M. B., Ruff, C. F., Yang, X., Ross, S. E. & Vazquez, A. L. Optogenetic assessment of VIP, PV, SOM and NOS inhibitory neuron activity and cerebral blood flow regulation in mouse somato-sensory cortex. J. Cereb. Blood Flow Metab. 40, (2020).

37. Uhlirova, H. et al. Cell type specificity of neurovascular coupling in cerebral cortex. Elife 5, (2016).

38. Anenberg, E., Chan, A. W., Xie, Y., LeDue, J. M. & Murphy, T. H. Optogenetic stimulation of GABA neurons can decrease local neuronal activity while increasing cortical blood flow. J. Cereb. Blood Flow Metab. 35, (2015).

39. Vazquez, A. L., Fukuda, M. & Kim, S. G. Inhibitory Neuron Activity Contributions to Hemodynamic Responses and Metabolic Load Examined Using an Inhibitory Optogenetic Mouse Model. Cereb. Cortex 28, (2018).

40. Devor, A. et al. Suppressed neuronal activity and concurrent arteriolar vasoconstriction may explain negative blood oxygenation level-dependent signal. J. Neurosci. 27, (2007).

41. Cauli, B. et al. Cortical GABA interneurons in neurovascular coupling: relays for subcortical vasoactive pathways. J. Neurosci. 24, (2004).

42. Lee, L. et al. Key Aspects of Neurovascular Control Mediated by Specific Populations of Inhibitory Cortical Interneurons. Cereb. Cortex 30, (2020).

43. Inoue, M. et al. Rational Engineering of XCaMPs, a Multicolor GECI Suite for In Vivo Imaging of Complex Brain Circuit Dynamics. Cell 177, (2019).

44. Han, S., Yang, W. & Yuste, R. Two-Color Volumetric Imaging of Neuronal Activity of Cortical Columns. Cell Rep. 27, (2019).

45. Lee, J. H. et al. Global and local fMRI signals driven by neurons defined optogenetically by type and wiring. Nature 465, 788–792 (2010).

46. Bernal-Casas, D., Lee, H. J., Weitz, A. J. & Lee, J. H. Studying Brain Circuit Function with Dynamic Causal Modeling for Optogenetic fMRI. Neuron 93, (2017).

47. Logothetis, N. K. What we can do and what we cannot do with fMRI. Nature 453, 869– 878 (2008).

48. Ekstrom, A. How and when the fMRI BOLD signal relates to underlying neural activity: the danger in dissociation. Brain Res. Rev. 62, (2010).

49. Howarth, C., Mishra, A. & Hall, C. N. More than just summed neuronal activity: how multiple cell types shape the BOLD response. Philos. Trans. R. Soc. Lond. B Biol. Sci. 376, (2021).

50. Gao, Y.-R. et al. Time to wake up: Studying neurovascular coupling and brain-wide circuit function in the un-anesthetized animal. Neuroimage 153, 382–398 (2017).

51. Han, Z. et al. Awake and behaving mouse fMRI during Go/No-Go task. Neuroimage 188, 733–742 (2019).

52. Madularu, D. et al. A non-invasive restraining system for awake mouse imaging. J. Neurosci. Methods 287, 53–57 (2017).

53. Zhang, W. et al. Acute stress alters the ‘default’ brain processing. Neuroimage 189, (2019).

54. Yoshida, K. et al. Physiological effects of a habituation procedure for functional MRI in awake mice using a cryogenic radiofrequency probe. J. Neurosci. Methods 274, (2016).

55. Hurst, J. L. & West, R. S. Taming anxiety in laboratory mice. Nat. Methods 7, (2010).

56. Power, J. D., Barnes, K. A., Snyder, A. Z., Schlaggar, B. L. & Petersen, S. E. Spurious but systematic correlations in functional connectivity MRI networks arise from subject motion. Neuroimage 59, (2012).

57. Ioanas, H. I., Marks, M., Zerbi, V., Yanik, M. F. & Rudin, M. An optimized registration workflow and standard geometric space for small animal brain imaging. Neuroimage 241, (2021).

58. Ioanas, H. I. et al. An Automated Open-Source Workflow for Standards-Compliant Integration of Small Animal Magnetic Resonance Imaging Data. Front. Neuroinform. 14, (2020).

59. Roche, A. A four-dimensional registration algorithm with application to joint correction of motion and slice timing in fMRI. IEEE Trans. Med. Imaging 30, (2011).

60. Avants, B. B. et al. A reproducible evaluation of ANTs similarity metric performance in brain image registration. Neuroimage 54, (2011).

61. Dorr, A. E., Lerch, J. P., Spring, S., Kabani, N. & Henkelman, R. M. High resolution three-dimensional brain atlas using an average magnetic resonance image of 40 adult C57Bl/6J mice. Neuroimage 42, (2008).

62. Jenkinson, M., Beckmann, C. F., Behrens, T. E., Woolrich, M. W. & Smith, S. M. FSL. Neuroimage 62, (2012).

63. Thévenaz, P., Ruttimann, U. E. & Unser, M. A pyramid approach to subpixel registration based on intensity. IEEE Trans. Image Process. 7, (1998).

64. Zhou, P. et al. Efficient and accurate extraction of in vivo calcium signals from microendoscopic video data. Elife 7, (2018).

65. Pnevmatikakis, E. A. et al. Simultaneous Denoising, Deconvolution, and Demixing of Calcium Imaging Data. Neuron 89, (2016).

