## Supplemental figures for "Simultaneous single-cell calcium imaging of neuronal population activity and brain-wide BOLD fMRI"

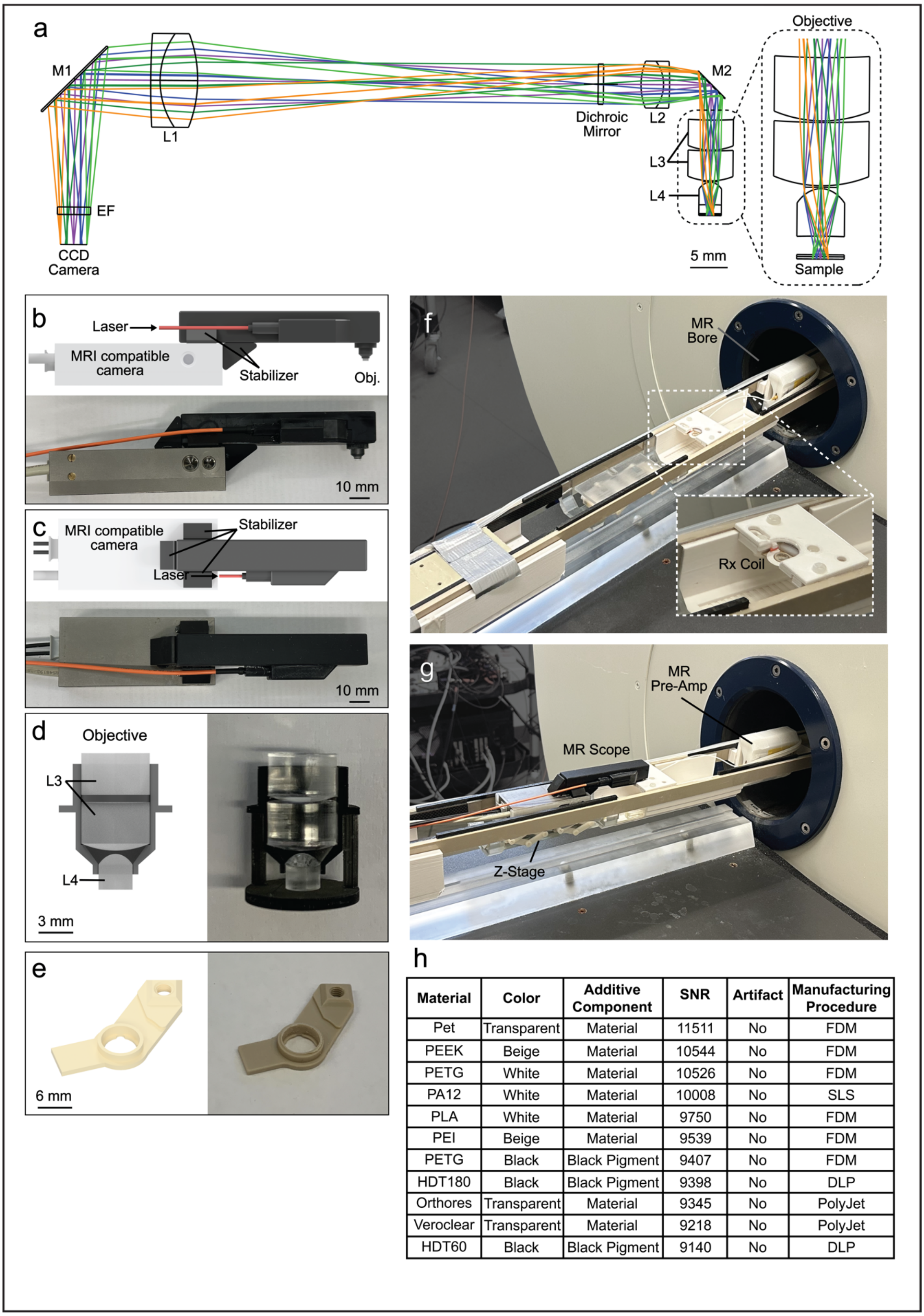


| Material | Color | Additive Component | SNR | Artifact | Manufacturing Procedure |
|---|---|---|---|---|---|
| Pet | Transparent | Material | 11511 | No | FDM |
| PEEK | Beige | Material | 10544 | No | FDM |
| PETG | White | Material | 10526 | No | FDM |
| PA12 | White | Material | 10008 | No | SLS |
| PLA | White | Material | 9750 | No | FDM |
| PEI | Beige | Material | 9539 | No | FDM |
| PETG | Black | Black Pigment | 9407 | No | FDM |
| HDT180 | Black | Black Pigment | 9398 | No | DLP |
| Orthores | Transparent | Material | 9345 | No | PolyJet |
| Veroclear | Transparent | Material | 9218 | No | PolyJet |
| HDT60 | Black | Black Pigment | 9140 | No | DLP |

**Supplementary figure 1 | MRI compatible microscope setup for combined BOLD fMRI and calcium imaging. a,** Zemax simulation of the emission optical pathway. Fluorescent emission is collected by an objective consisting of three small diameter acrylic aspheric lenses (L3, L4). The collected light is relayed through two achromatic lenses (L1, L2), and projected onto the MRI compatible CCD camera through an emission filter (EF). **b,** Side view of the MRI compatible microscope. **c,** Top view of the MRI compatible microscope. **d,** Cross-section of the objective consisting of three small diameter acrylic aspheric lenses (L3, L4). The real image (right) includes the support structure used for aligning the bottom lens (L4) during construction. **e,** The custom designed PEEK headbar. **f,g,** Picture of the complete imaging setup including the microscope, translational stage for focusing, and the MRI scanner. The inset in **f** shows the head fixation stage, and receiver (Rx) coil. **h,** Table showing the BOLD fMRI signal to noise (SNR) values of tested materials. Higher SNR values correspond to better quality BOLD fMRI images.

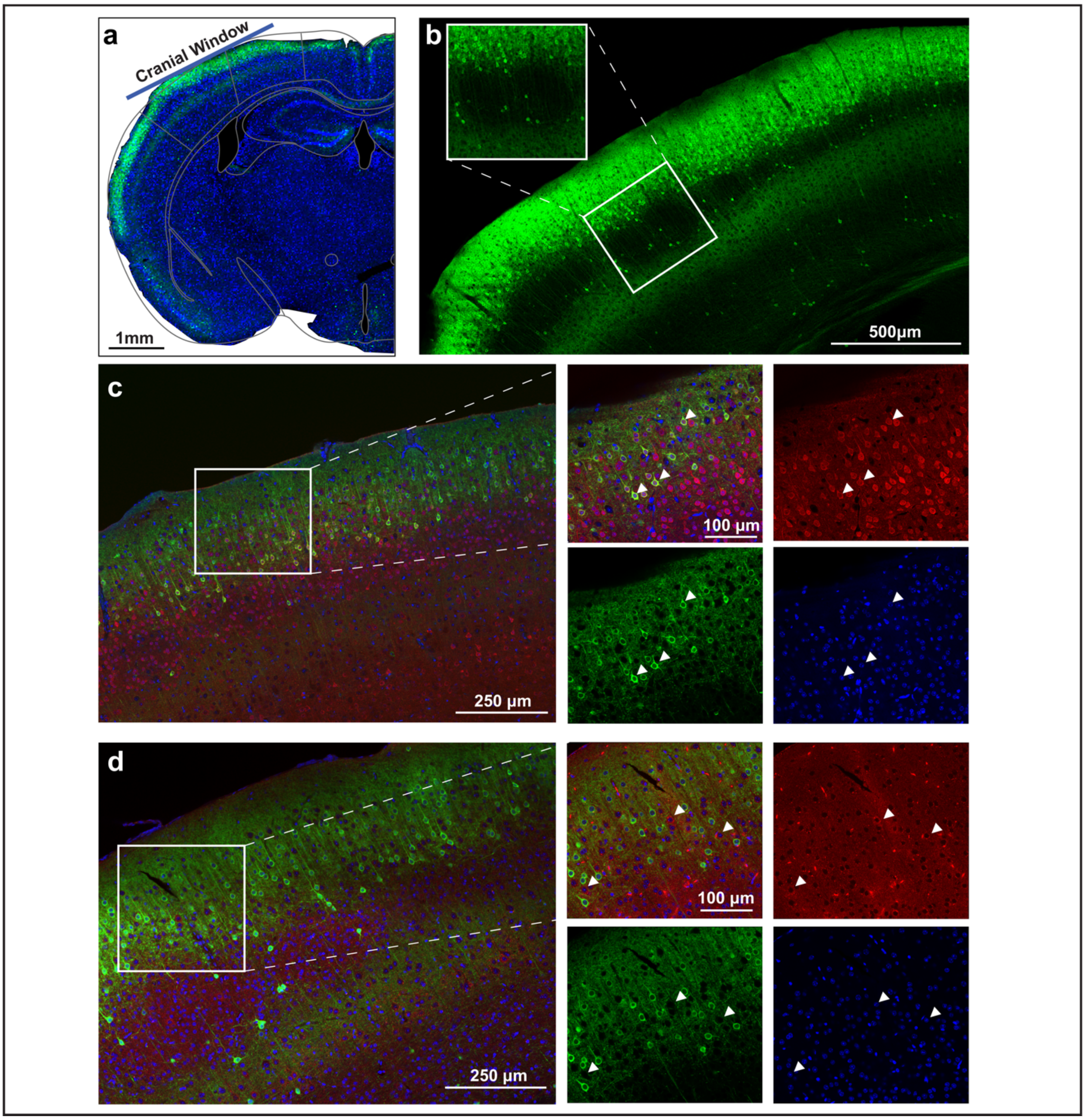


**Supplementary figure 2 | GCaMP-labeled cell-type verification. a,** Coronal brain slice stained with DAPI (blue) and expression of GCaMP6M (green) in layer 2/3 of a Rasgrf-2A-Cre animal. The cranial window is shown as a blue line located above the SSp-bfd. **b,** Close-up of the same slice with a blow-up of the barrel fields. **c,** Immunohistological verification of different cell-types. Neurogranin positive cells are shown in red, GCaMP is shown in green, and DAPI in blue. **d,** Immunohistological verification of different cell-types. GAD65+67 positive cells are shown in red, GCaMP is shown in green, and DAPI in blue.

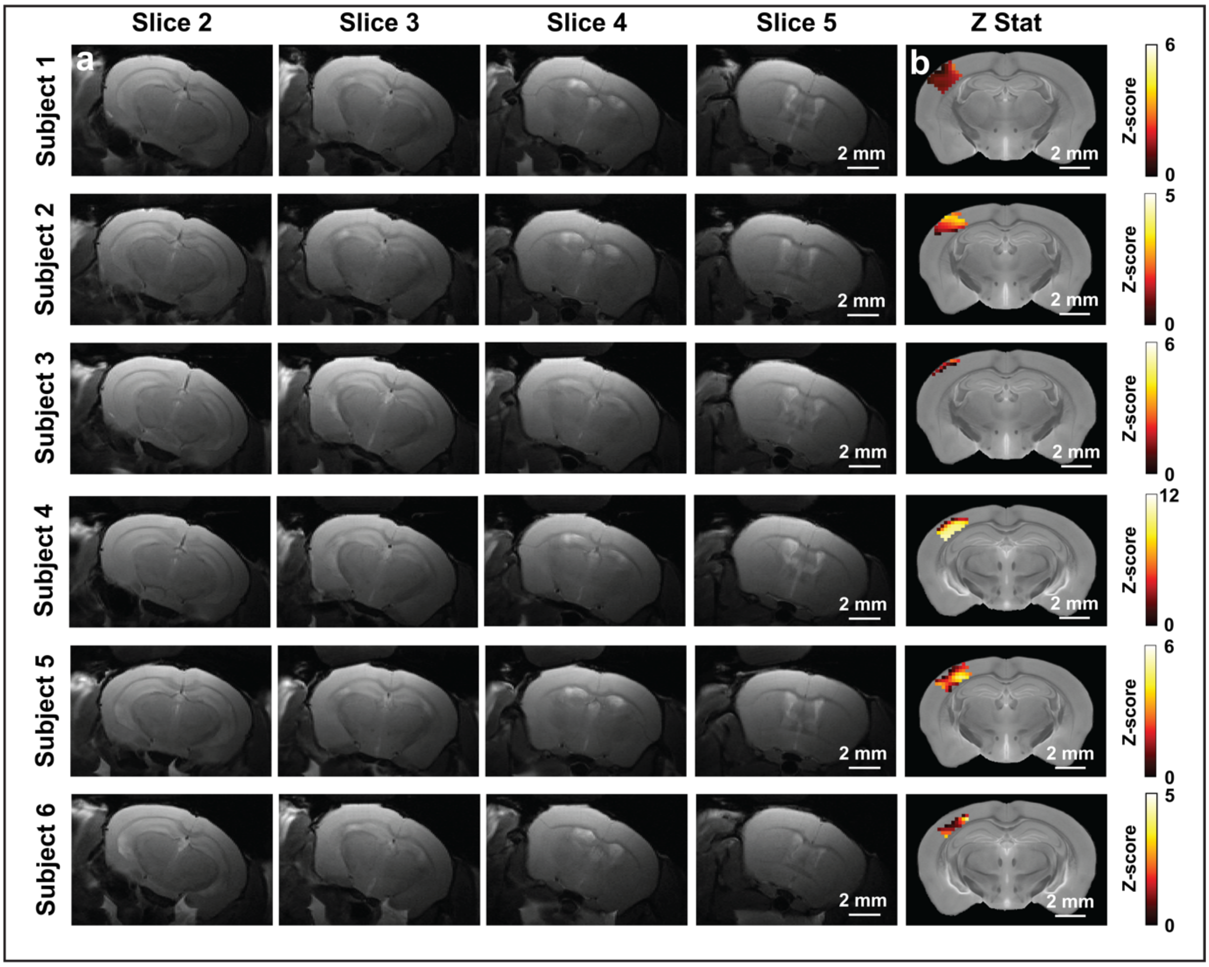


**Supplementary figure 3 | Coronal MRI slices around the glass-window implant used for optical imaging, and General Linear Model based statistical maps for individual animals. a,** Unprocessed anatomical images demonstrate the quality of the surgical procedure, and do not show any residual blood or air-bubbles. In addition, the image does not show any significant artifacts due to the implant. **b,** The GLM statistical parametric map for each animal (z-score ≥ 3.1, FWE *P* ≤ 0.000001) shows significantly activated voxels in the SSp-bfd for each mouse.

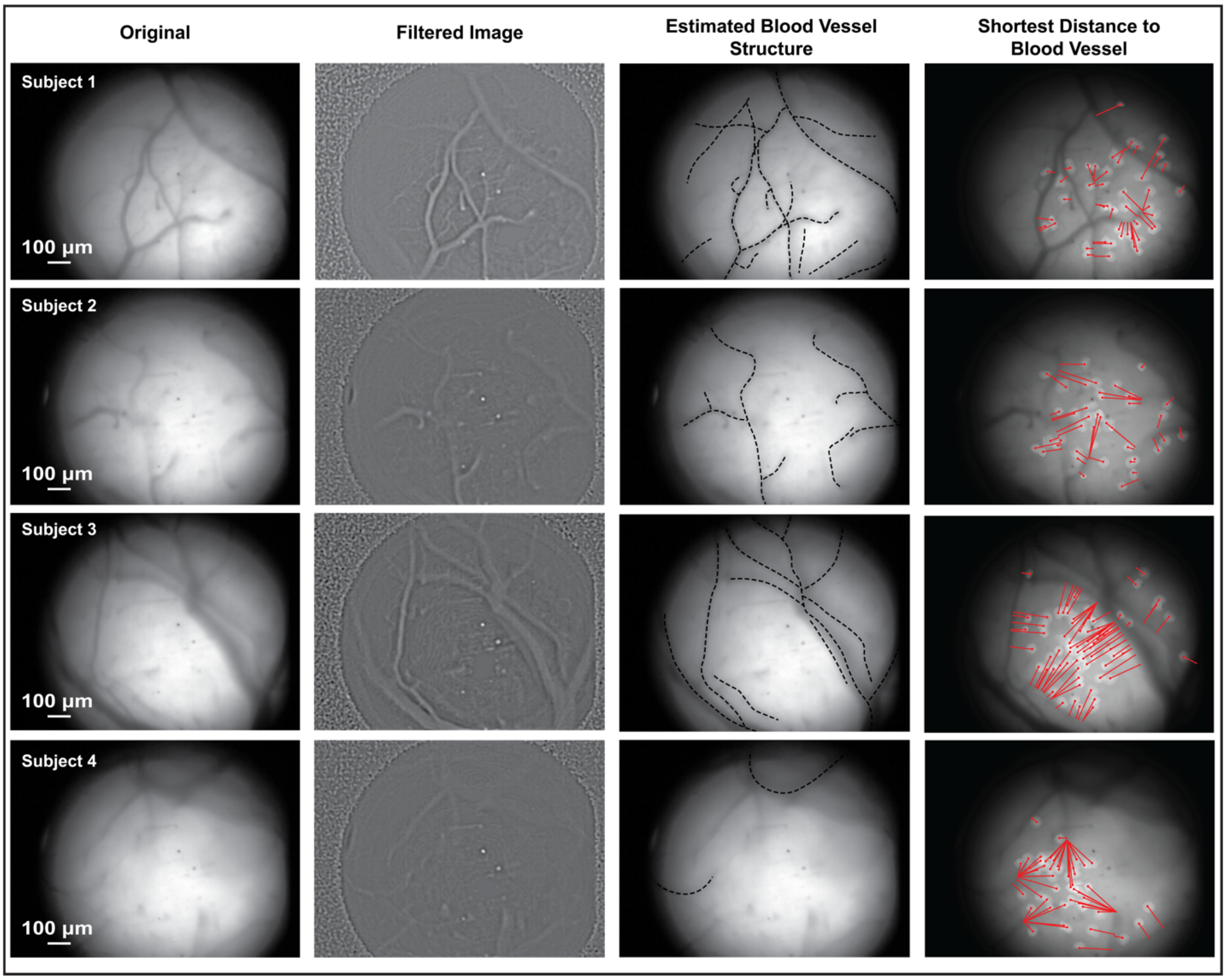


**Supplementary figure 4 | Annotation of the blood vessel patterns for individual animals.** The neural-vascular proximity was estimated based on the manual annotation of the vessel structure in the region of interest for each mouse. We used the time-averaged microscopic field (first column) and the band-passed filtered (100-150 Hz) microscopic field (second column) to accurately distinguish the vessel structure (third column). The neural-vascular proximity was calculated as the minimal Euclidean distance between the center of the cell and the vascular structure (fourth column).

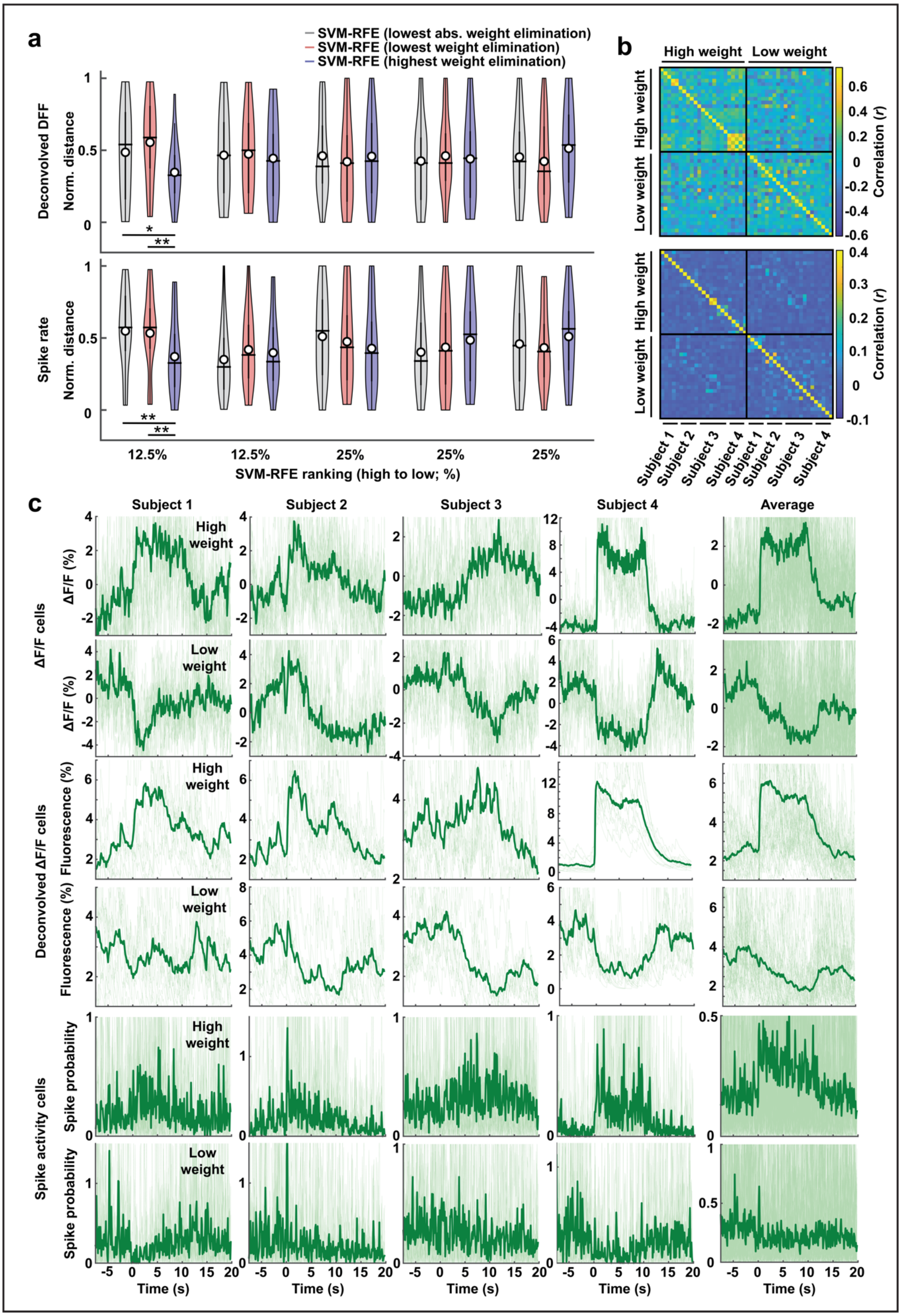


**Supplementary figure 5 | The effect of neuron-vascular proximity on the predictive relationship between BOLD fMRI and single-cell calcium activity is independent of the**

**ΔF/F calcium activity processing steps. a,** The neuron-vascular proximity measures, quantized based on the SVM decoding procedure in **Fig. 3a**. Features were ranked using recurrent feature elimination (see methods) using three individual ranking criteria (highest weights elimination, lowest weights elimination, and lowest absolute weight elimination). In contrast to **Fig. 3e**, we used either the deconvolved ΔF/F $Ca^{2+}$ activity or the spike probability estimate extracted using the CNMF-e pipeline (deconvolution = constrained FOOPSI algorithm, decay_time = 0.4; 64) as input for the SVM-RFE. Results for each extraction method are similar to those presented in **Fig. 3**. The central line represents the median, the circle represents the mean. The bold whisker extends to the 25th and 75th percentiles, the fine whisker extends to all data points. The width of the plot is proportional to the number of data points with the corresponding value. We tested for the difference between each quantile using a two-sample *t*-test (two-sample *t*-test; * = $P \leq 0.1$; ** = $P \leq 0.05$). No correction for multiple comparisons was applied. **b,** The correlation matrix between the ΔF/F $Ca^{2+}$ activity of the highest ranked neurons based on the ranking criteria (highest weights elimination, lowest weights elimination). **c,** Trial average neuronal $Ca^{2+}$ activity of the highest ranked cells for two ranking criteria (lowest weights elimination, top; highest weights elimination, bottom). Data is presented for the ΔF/F $Ca^{2+}$ activity, deconvolved ΔF/F $Ca^{2+}$ activity, and the spike probability (deconvolution = constrained FOOPSI algorithm, decay_time = 0.4; 26, 65). Results are similar across different extraction methods.

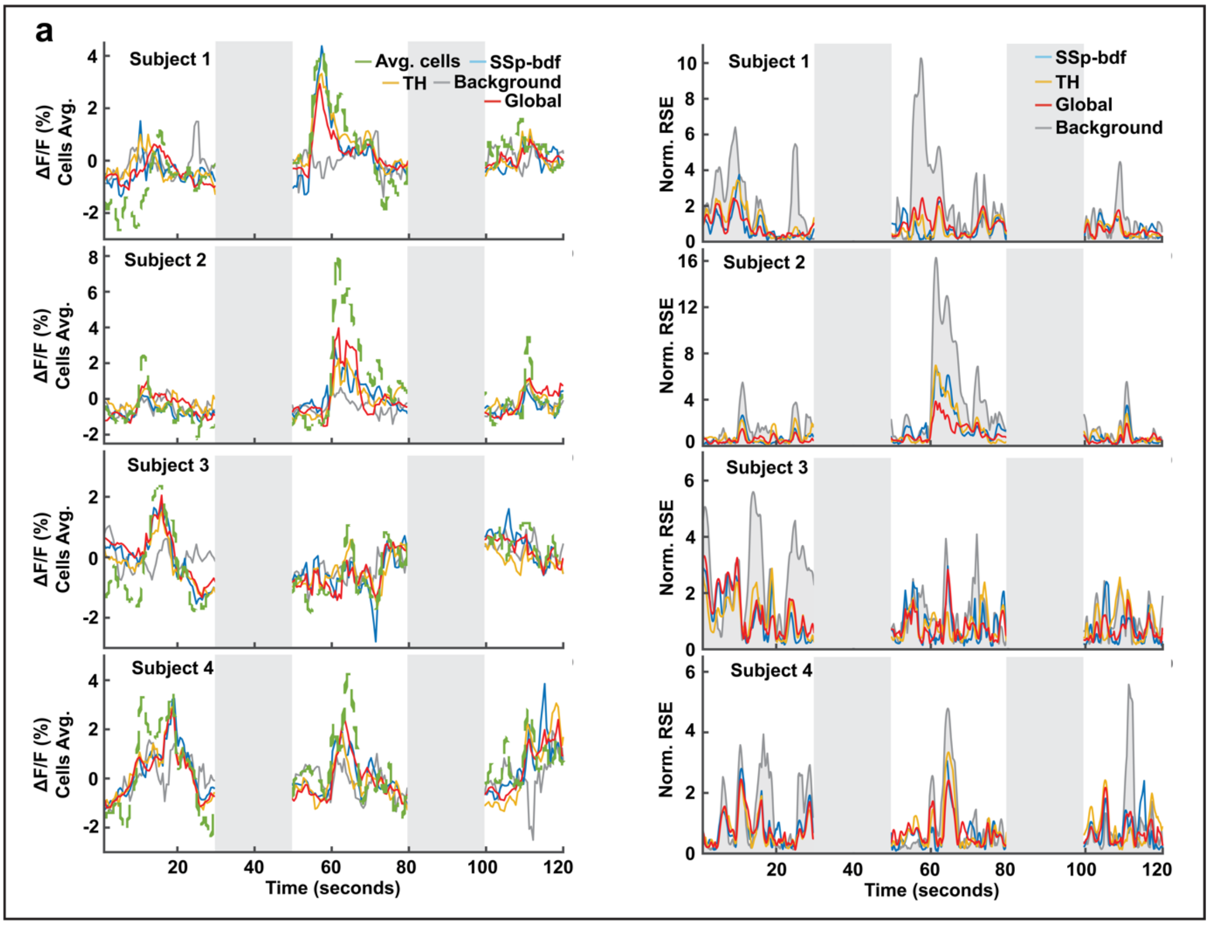


**Supplementary figure 6 | Prediction of the BOLD fMRI from the SSp-bfd ΔF/F population activity.** Example predictions of the SSp-bfd ΔF/F population fluorescent calcium activity (left) and corresponding normalized root square error (right; NRSE) for different brain areas and background noise. Results are plotted for individual mice. Gray boxes indicate laser-off period. SSp-bfd = somatosensory barrel fields, TH = thalamus.
